# Spatio-temporal coordination of virulence, metabolism, and stress responses shapes infection dynamics of *Xanthomonas perforans*

**DOI:** 10.64898/2026.01.29.702562

**Authors:** Amanpreet Kaur, Sivakumar Ramamoorthy, Palash Ghosh, Kylie Weis, Neha Potnis

**Author notes:** These are co-first authors (Amanpreet Kaur, Sivakumar Ramamoorthy). These authors have equal contribution (Palash Ghosh, Kylie Weis). Corresponding author: Neha Potnis.

## Abstract

Plants provide distinct ecological niches for diverse microbial communities, with each member adopting strategies tailored to the specific ecological niche it inhabits. Two foliar niches, the leaf surface (epiphytic environment) and the apoplast, impose distinct physiological constraints on microbial fitness, particularly for hemibiotrophic pathogens. In this study, we investigated how these environments shape the transcriptional responses of *Xanthomonas perforans* (*Xp*), a tomato pathogen, and how its virulence factors, metabolic pathways, and regulatory networks are spatially and temporally coordinated during disease progression. Transcriptome profiling of a pathogen recovered from the leaf surface and apoplast revealed pronounced niche-specific and colonization stage-specific gene expression patterns. Early epiphytic colonization was characterized by activation of chemosensing, and motility pathways that facilitate pathogen relocation and acquisition of limiting nutrients such as iron and phosphate. This stage also featured induction of DNA and protein repair systems, quorum sensing pathways, phenylalanine degradation and tyrosine conversion to counter phenylpropanoid defenses, genes involved in mitigating osmotic and oxidative stress, active DNA exchange machinery, and type VI secretion system-mediated microbial competition. Upon entry into the apoplast, *Xp* shifted toward active metabolism and replication, accompanied by investment in type II and III secreted virulence factor expression. Genes involved in evasion of plant immunity and overcoming of host-mediated nutrient sequestration were also upregulated, including those involved in quinone detoxification, phosphate and sulfur uptake, and fatty acid, xanthan, and LPS biosynthesis. During late apoplastic colonization, the pathogen transitioned again towards strong stress response activation, followed by renewed expression of flagellar motility and chemotaxis genes, suggesting preparation for dissemination. Notably, genes associated with oxidative and nutrient stress were enriched across both niches, although specific components differed. Type IV pili, conjugation genes, and plasmid-borne type III effectors were induced early in both niches, suggesting their niche-independent role in initial establishment. Together, these findings reveal a coordinated spatio-temporal regulatory strategy during the transition from the leaf surface to the apoplast.

**Author Summary:** *Xanthomonas perforans* is a foliar bacterial pathogen that infects tomato plants and leads to severe yield losses. To establish a successful infection, the pathogen must overcome a series of environmental and host-imposed challenges. This study characterizes the traits activated at distinct stages of infection, during both early and late pathogenesis, and across different niches, including the leaf surface and its interior (apoplastic) space. On the leaf surface *Xanthomonas* mainly focuses on movement, communication, and survival against stress and starvation with the major functions related to motility, nutrient uptake, and DNA transfer during early stages. Once inside the leaf, the bacteria switches tactics to focus primarily on reproduction, defense against the plant immune response, production of factors that weaken the plant’s defenses and gaining access to nutrients the plant normally restricts. Understanding the different stages of infection may inform how the crosstalk among host and pathogen unfolds during pathogenesis allowing us to understand the host environment. These findings can help us discover pathogen weaknesses that could be targeted for disease management.

## Introduction

Plants present multi-compartment niches for diverse microbes, including both beneficial and detrimental ones that influence growth and overall productivity. Each niche is shaped by distinct physical, chemical, and biological constraints [1–3]. For foliar microbes, leaf surfaces and the extracellular space inside plant tissue (apoplastic space) represent two ecological niches. The leaf surface is vast, nutritionally poor, and environmentally variable with exposure to UV radiation, fluctuating humidity and temperature, episodic desiccation; and microbial competition for space and resources [4]. On the other hand, the apoplast shields microbes from UV radiation and desiccation, offering opportunities to exploit host resources for multiplication while simultaneously challenging the pathogens with immune defenses and antimicrobial compounds [3]. On leaf surfaces, bacteria contend with nutrient and water limited conditions, where even simple carbon sources and amino acids can be spatially patchy and diffusion limited [5]. To locate these nutrient oases and withstand environmental stresses, bacteria rely on motility, chemotaxis, and biofilm formation [6–10]. In addition, coordinated polysaccharide production, osmo-tolerance and stress response programs further enhance bacterial survival under these constraints [11–13]. Epiphytic persistence is not merely passive but is a critical determinant to disease outcome, as surface population size is correlated with host entry and subsequent symptom development [14,15]. Entry into leaves typically occurs through natural openings such as stomata or hydathodes. Stomata actively participate in the innate immune response, which pathogen can overcome by deploying toxins and effectors to circumvent stomatal closure and gain access to the apoplast [16–18]. Once inside the apoplast, the pathogen population expands, and disease is often associated with sites harboring large internal bacterial populations [19]. Water availability becomes a central bottleneck in the apoplast. Successful pathogens manipulate host processes to transiently hydrate the apoplast (water-soaking), thereby facilitating bacterial proliferation. In *Pseudomonas syringae*, effectors, AvrE1 and HopM1 are known to induce apoplastic water-soaking [11,20]. In *Xanthomonas*, Transcription-Activator-Like effector, AvrHah1, was shown to induce water-soaking and promote proliferation [21].

Because leaf surface and apoplast impose fundamentally different constraints, successful foliar pathogens must reprogram gene expression across space and time. In *P. syringae*, epiphytic stages emphasize active relocation in response to nutrients and water, whereas apoplastic growth favors metabolic reconfiguration, secretion system deployment, and stress tolerance [11,15,22]. Early *in planta* transcriptomics further shows that plant immunity reshapes pathogen iron acquisition, secretion systems, and stress responses, linking bacterial gene expression states to later disease outcomes [23]. Similar niche-dependent patterns were observed in *Xanthomonas campestris* pv. campestris during hydathode infection [26], including repression of motility and induction of sulfur and phosphate metabolism. These studies revealed that the driving forces for pathogen adaptation to epiphytic and apoplastic environments are distinct, emphasizing the importance of capturing the gene expression across both spatial and temporal stages.

The genus *Xanthomonas* comprises a diverse group of Gram-negative bacteria responsible for causing disease in over 400 plant species and yield losses of major economically important crops [27]. Although individual virulence factors, primarily, secretion systems (type I to type VI) and their associated effectors, along with exopolysaccharides and quorum sensing systems have been well characterized in *Xanthomonas* [27,28], how their deployment is coordinated across space and time during infection is far less clear. *Xanthomonas perforans* (*Xp*) is also a hemibiotrophic seedborne pathogen within the genus *Xanthomonas*, and the causative agent of bacterial leaf spot (BLS) disease in tomato and pepper. *Xp* has a pronounced epiphytic phase allowing the pathogen to establish on the leaf surface before entering the biotrophic phase [29]. To elucidate the spatio-temporal adaptive strategies of *Xanthomonas perforans* during tomato colonization, we conducted genome-wide transcriptome profiling of *Xp* strain AL65 across spatial niches of leaf surface (at 8 hours and 24 hours after dip-inoculation) and apoplastic space (24 hours and 72 hours after infiltration), with three biological replicates per time point **(Fig 1)**. This spatio-temporal design enabled us to capture transcriptional reprogramming associated with early and late stages of infection. We leveraged Weighted Gene Co-expression Network Analysis (WGCNA) to define niche- and time-specific transcriptional programs employed by *Xp* during tomato colonization, to probe the host environment perceived by the pathogen, and to understand how the pathogen modulates the host environment during the infection process. We centered our interpretation on biologically informative genes within each co-expression module. Key genes were organized into functional categories and their differential expression was corroborated, enabling integrated interpretation of co-expression patterns and expression levels.

**Fig 1.**
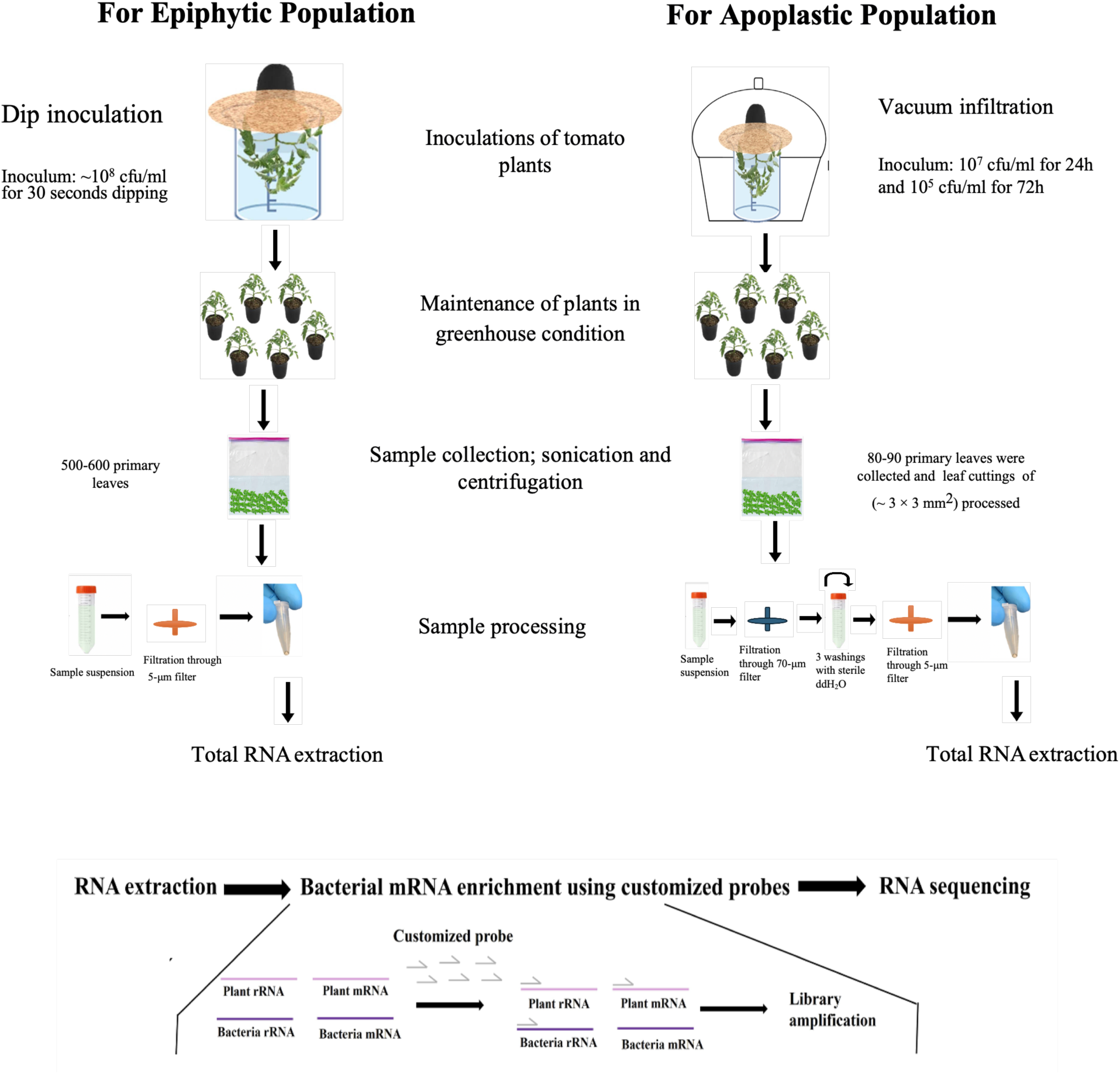
The figure illustrates the experimental design used to establish bacterial populations in epiphytic and apoplastic niches. Epiphytic populations were generated by dip inoculation with an inoculum concentration of ∼10⁸ CFU/mL for both 8 hours and 24 hours sampling points. In contrast, apoplastic populations were established by vacuum infiltration using initial concentrations of 10⁷ and 10⁵ CFU/mL for sampling at 24 hours and 72 hours post inoculation timepoints respectively. Sampling was performed and followed by filtration to remove plant material. For epiphytic samples, a 5-µm filter was used, whereas apoplastic samples were first passed through a 70-µm filter and then a 5-µm filter to remove host tissue from leaf cuttings and sonicated sample material. Total RNA was then extracted, and bacterial mRNA was enriched using host-specific depletion probes, which are listed in Supplementary Table 1.

## Materials and Methods

### Plant material, bacterial strain and growth conditions

The tomato cultivar FL-47 was used in this study. Two-week-old seedlings were transplanted into 4-inch pots with soil less standard premix potting medium. Plants were grown at 28 to 30°C under greenhouse conditions. For the experiment, 4 to 5 weeks old plants were used. The *Xanthomonas perforans* (*Xp*) strain used in this experiment was AL65 (GCF_007714115.1), originally isolated from pepper (*Capsicum annuum*) in Alabama, USA [30]. The strain was routinely grown in nutrient medium at 28°C with or without agitation.

### *In planta* bacterial growth for RNA extraction

The AL65 strain was streaked on nutrient agar plates and grown for 36 hours at 28°C. The cells were scraped off the plates using sterile 0.01M MgSO_4_ for inoculum preparation. The cells were washed two times using sterile 0.01M MgSO_4_. The workflow for *in planta* transcriptome is shown in **Fig 1**. To establish an epiphytic population the inoculum density was adjusted to ∼10^8^ cfu/ml (OD_600nm_ = 0.3) and amended with 0.01M MgSO_4_ containing 0.025% Silwet and the plants were dipped into the inoculum for 30 seconds such that the abaxial and adaxial surface of the leaves were completely immersed in the inoculum. The dip-inoculated plants were maintained in greenhouse conditions and leaves were collected at 8 hours (8h) and 24 hour (24h) post-infection. A total of 500-600 primary leaves were collected, immediately submerged in 2 L RNA stabilization solution (10 mL water-saturated phenol (pH <7.0), 190 mL ethanol, 1.8 L water), sonicated for 10 minutes and centrifuged at 5,000 × g for 10 min to collect bacterial cells. The cell pellets were suspended in residual supernatant and filtered through a 5-μm filter (Millex-SV syringe filter unit; Millipore Corp.). The cells in the filtrate were harvested by centrifugation at 5,000 × g for 5 min, the supernatant was discarded, and the pellets were placed at −20 °C. The cells collected from the 500–600 leaflet from 50 plants per replicate at a time point served as a biological replicate, and the procedure was repeated to provide three biological replicates for each time point.

To establish apoplastic populations, bacteria were introduced into the plant by vacuum infiltrating leaves submerged in the inoculum. The plants and bacteria were grown, and the inoculum was prepared, as described above. We used a different inoculum density for each time point, 10^7^ cfu/ml for 24h timepoint to allow recovery of enough cells during early time point of 24h post-inoculation and 10^5^ cfu/ml for the 72 hour (72h) timepoint to capture pathogen gene expression during later stages of disease development while avoiding necrosis of the plants before the 72h sampling point. Infiltrated plants were incubated for 24h and 72h in greenhouse conditions. A total of 80-100 leaves were collected and submerged immediately in an acidic phenol RNA-stabilizing solution as described above. The leaves were cut into squares (∼3 × 3 mm^2^) while submerged, and the plant tissues and liquid were sonicated for 10 min. The solution was filtered through 70-µm filter with repeated filter changes to remove the host contamination or debris. The filtrate was centrifuged at 7,000 × *g* for 10 min, and the pellet was suspended in the remaining supernatant, which was subjected further to three washings with sterile ddH_2_O to eliminate plant material. The suspension was filtered through a 5-µm filter with repeated filter changes. The bacterial cells were harvested by centrifugation at 7,000 × *g* for 10 min, the supernatant was discarded, and the pellets were flash frozen and placed at −20 °C. The cells collected from the 80-100 leaves on a single time point served as a biological replicate, and three biological replicates were generated at each time point.

### RNA extraction, ribodepletion and sequencing

Total RNA was extracted using TRIzol (Thermo Fisher Scientific, Waltham, MA, USA) as described previously [31]. The concentration of RNA samples was determined using QubitRNA HS Assay kit (Invitrogen). Ribodepletion was performed using customized probes targeting the *Xp* AL65 rRNA and for removal of plant derived mRNA and rRNA probes were designed targeting tomato, which are listed in **S1 Table**. The enriched bacterial mRNA was converted to cDNA, and the sequencing library was generated.

### Analysis of RNA-Seq results

The sequenced data was processed to remove adapters and the FASTQ files were generated. We used the epinatel pipeline (https://github.com/epinatel/Bacterial_RNAseq/). The pipeline uses steps involving read quality filtering using trimmomatic [32], mapping to the reference genome of *Xp* AL65 (RefSeq: NZ_SMVI00000000.1, assembly: GCF_007714115.1) using bowtie2 [33]. Differentially expressed genes (DEGs) across the four treatment conditions (apoplastic and epiphytic each with two time points) were identified using the DESeq2 R package (v1.46.0) [34].This method analyzes RNA-seq count data using a model based on the negative binomial distribution to detect differences in gene expression between treatments. The resulting *p*-values were adjusted for multiple testing using the Benjamini–Hochberg method to control the false discovery rate (FDR). Genes with an adjusted *p*-value of less than 0.05 were considered significantly differentially expressed and were used for further functional enrichment analysis. Gene ontology analysis was performed using EggNOG (v.5.0) [35]. KEGG pathway enrichment analysis was performed on the differentially expressed genes using the clusterProfiler R package (v4.14.6). This analysis was used to identify biological pathways that were significantly overrepresented among the differentially expressed genes. KEGG pathways with an adjusted *p*-value of less than 0.05 were considered significantly enriched (http://www.genome.jp/kegg/).

### Gene co-expression network analysis

Gene co-expression network analysis was performed using RStudio with the WGCNA package (v1.73) [36]. Gene read counts were filtered to retain genes with ≥10 reads in at least three samples and then normalized using the variance stabilizing transformation from the DESeq2 package (v1.49.4). The normalization accounted for differences in sequencing depth across apoplast vs epiphytic conditions. To construct the co-expression network, a soft-threshold power (β = 7) was chosen, which was the first value to reach a correlation coefficient of 0.82, for computing the adjacency matrix (measuring connection strength between genes). This matrix was then used to calculate the topological overlap measure (TOM), using the “signed” option to ensure modules capture biologically meaningful patterns where genes exhibit coordinated up- or down-regulation. Pairwise gene distances derived from TOM were used to perform hierarchical clustering, and modules were initially identified using the cutreeDynamic function with deepSplit = 2 and a minimum module size of 30. Initially, 17 modules were detected, which were subsequently merged based on ≥75% similarity (MEDissThres = 0.25) using the mergeCloseModules function, resulting in a final set of 8 co-expression modules. Module–trait relationships were assessed using Pearson’s correlation. Top genes within each module were selected based on a module membership (kME) value greater than 0.8, and correlations with *p-*value < 0.05 were considered statistically significant.

## Results and Discussion

### *Xp* demonstrates niche-specific expression patterns during colonization and proliferation in epiphytic and apoplastic environment

To identify coordinated gene expression patterns, we applied Weighted Gene Co-expression Network Analysis (WGCNA), which clusters genes into modules based on highly correlated expression profiles. This approach revealed seven distinct modules (**Fig 2**), each representing functionally related gene sets. Modules were prioritized for interpretation based on strong correlations with specific infection stages and niches. After filtering for high module membership (kME ≥ 0.8, *p*-value < 0.05), we identified 311 genes in the black module, 509 in blue, 514 in green, 141 in greenyellow, 341 in magenta, 120 in midnight blue, 136 in purple, and 289 in yellow. Notably, the green and yellow modules were niche-specific, corresponding to epiphytic and apoplastic conditions respectively. Within the apoplast, the blue and midnightblue modules were associated with early infection (24h), whereas the black module represented late infection (72h). No temporal separation was evident during epiphytic colonization. The purple module marked early colonization in both niches, whereas the magenta module represented overlapping expression during late apoplastic and epiphytic colonization. Differential expression analysis using DESeq2 in parallel corroborated module level trends observed in WGCNA. With DESeq2, we observed 920 genes were upregulated and 943 genes were downregulated in epiphytic conditions compared with the apoplast. Further, 169 genes were upregulated and 162 were downregulated at 8h compared with 24h in the epiphytic niche, whereas 816 genes were upregulated and 786 were downregulated at 72h compared with 24h in the apoplast niche (**Fig 3**). To gain insight into the biological processes associated with these transcriptional changes, KEGG pathway enrichment analysis was performed on the DEGs, and the significantly enriched pathways are summarized in **Fig 3** (**S2 Table**).

**Fig 2.**
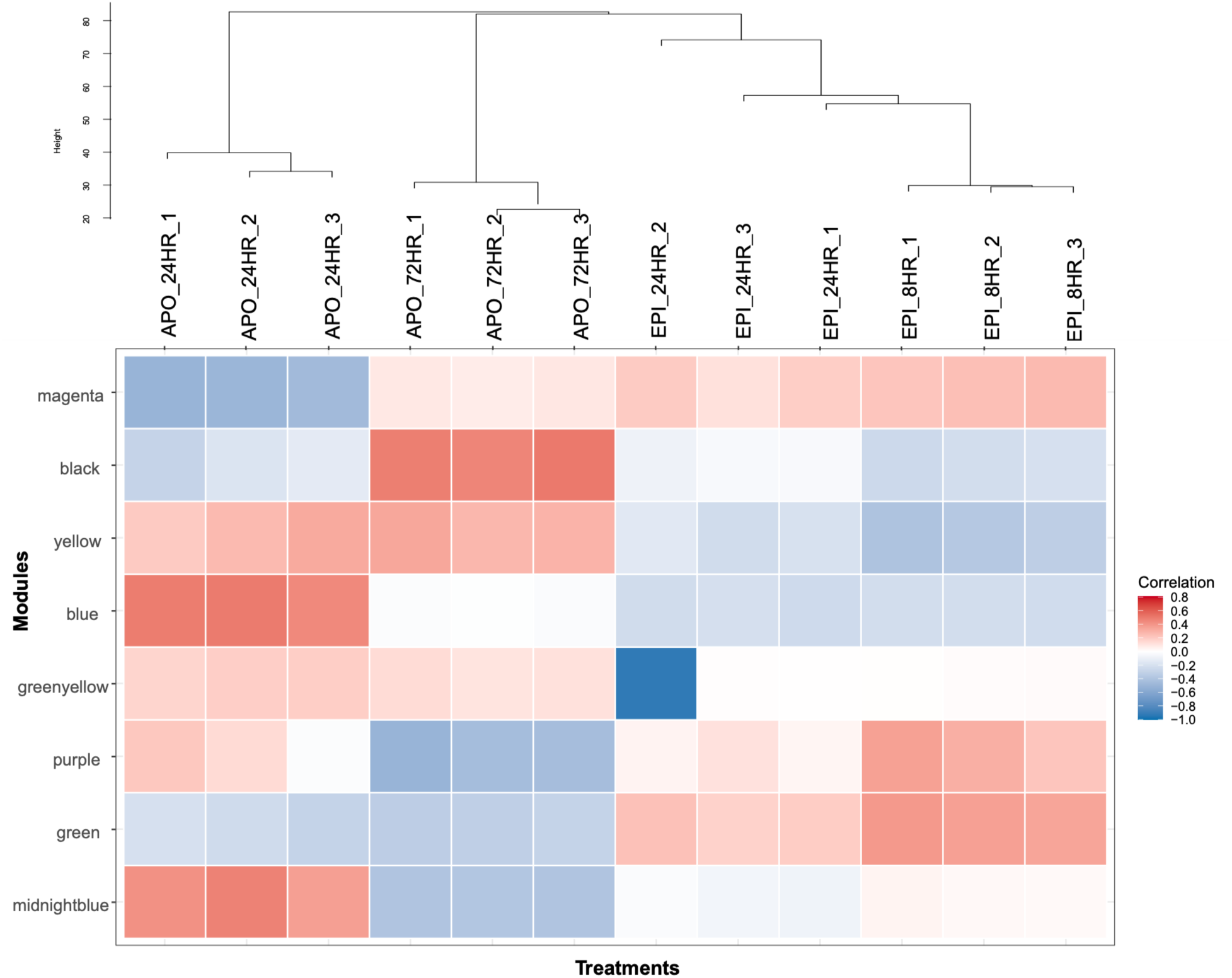
The heatmap illustrates the modules identified by Weighted Gene Co-expression Network Analysis (WGCNA), which group genes based on pairwise expression distances. The y-axis shows the different modules (blue, green, yellow, purple, magenta, midnight blue, greenyellow, and black), while the x-axis represents the samples arranged with dendrogram. The color scale indicates the correlation between each sample and each module: darker red corresponds to higher correlation, implying stronger expression of the module genes in those samples, whereas lighter colors indicate weaker associations. *(Note: the name tags of x-axis (also on dendrogram tip) are referring to samples, where APO means Apolastic, and EPI means Epiphytic samples with 1,2,3 as replicate numbers for each specific treatment with timepoints.)*

**Fig 3.**
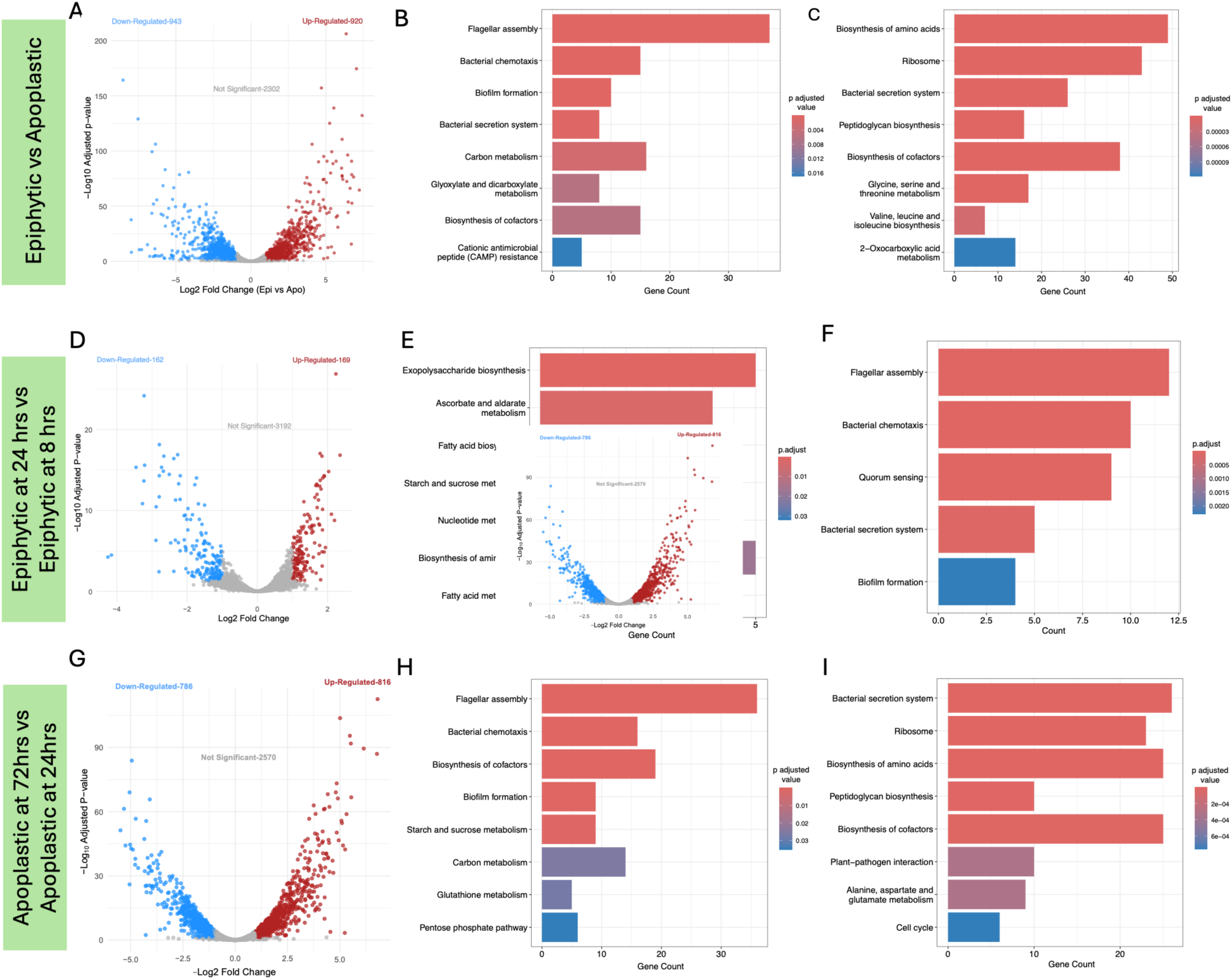
Transcriptomic analysis of *Xanthomonas perforans (Xp)* during colonization of epiphytic and apoplastic niches in tomato plants. (A, D, G) Volcano plots showing differentially expressed genes (DEGs) of *Xp* under the following comparisons: epiphytic vs apoplastic niches; epiphytic at 24 hours vs epiphytic at 8 hours; and apoplastic at 72 hours vs apoplastic at 24 hours, respectively. Up- and down-regulated genes are indicated by red and blue dots, respectively. (B, E, H) KEGG pathway enrichment analysis of up-regulated DEGs for the corresponding comparisons, with color indicating the adjusted *p*-value. (C, F, I) KEGG pathway enrichment analysis of down-regulated DEGs for the same comparisons, with color indicating the adjusted *p*-value.

Here we focused on WGCNA modules to define niche- and time-specific transcriptional programs employed by *Xp* during tomato colonization. All genes significantly associated with each module are listed in **S3 Table**. The key genes within each module were further functionally categorized and their expression levels were confirmed using DESeq2 log_2_ Fold-Change (log_2_FC) values (**S4 Table**).

### Traits specific to the leaf surface niche include chemotaxis, quorum sensing, motility, conjugal transfer, cellulose synthesis, active nutrient acquisition, DNA and protein repair, metabolism, and mechanisms for stress tolerance in *Xp*

The epiphytic niche was defined by the green module (514 genes), with 440 genes significantly upregulated relative to the apoplast (adjusted *p* < 0.05). Temporal differentiation was minimal, as only 157 genes were upregulated, and one gene downregulated at 8h compared with 24h, indicating largely stable epiphytic expression across timepoints.

#### Chemosensing, chemotaxis, motility, and quorum sensing

Of the 48 methyl-accepting chemotaxis proteins (MCPs) encoded in the genome, 22 were assigned to the epiphytic-specific green module and were strongly upregulated relative to the apoplast (log_2_FC: 1-5). MCPs showing the highest induction (log_2_FC: 4-5) were frequently adjacent to genes encoding hypothetical proteins. Notably, two MCPs, E2P69_RS20880 (log_2_FC: 2.911) and E2P69_RS20895 (log_2_FC: 3.708), flank the flagellar brake gene, *ycgR* (log_2_FC: 3.156). YcgR functions as a c-di-GMP-responsive “backstop brake” that modulates flagellar motor direction and speed, enabling fine-tuning of swimming velocity in response to environmental cues [37,38]. This genomic organization suggests MCP-guided regulation of flagellar motility under epiphytic growth. Consistent with this inference, multiple flagellar genes, including *motB2* (E2P69_RS04570), *motA* (E2P69_RS04575), and *fliJ* (E2P69_RS17055) were represented in the green module and were upregulated under epiphytic conditions (log_2_FC: 3-4). However, average flagellar transcript levels declined at 24h of epiphytic growth (avg. log_2_FC: –1.1), indicating a progressive shift away from active swimming as surface colonization proceeds. MotA and MotB form the stator complex driving flagellar rotation, FliJ contributes to flagellar assembly and virulence [39,40].

Genes involved in type IV pili biogenesis (*pilGHIJTUMNAWT*) were strongly upregulated at 8h, together with the mechanotaxis-associated genes, *pilJ* and *pilI*. These genes were co-expressed with the twitching motility response regulators *pilG* (log_2_FC: 1.66) and *pilH* (which negatively regulates swimming motility and positively regulates swarming motility), indicating coordinated regulation of surface-associated and collective motility during early epiphytic colonization. While most chemotaxis and chemosensing genes were significantly more highly expressed at 8h than at 24h, *pilJ* and *pilI*, showed no temporal differences, suggesting sustained mechanosensory signaling.

Quorum sensing and cyclic-di-GMP signaling were also prominently regulated under epiphytic conditions. The entire *rpf* operon clustered in the green module was significantly upregulated relative to the apoplast, indicating a coordinated regulatory response. The *rpfF* gene was upregulated at 8h (log_2_FC: 0.857), suggesting early DSF production, whereas both *rpfC* and *rpfG* genes, were consistently upregulated (log_2_FC: 0.922 and 0.798, respectively), with no significant temporal differences, indicating sustained DSF sensing, and c-di-GMP turnover promoting transition from a sessile to a motile state, and modulating biofilm formation. Co-expression of three GGDEF-domain diguanylate cyclases, two EAL-domain phosphodiesterases, one HD-GYP-domain protein indicates fine-scale modulation of c-di-GMP homeostasis. Additional regulator linked to cyclic-di-GMP signaling, transcriptional activator Zur was significantly upregulated under epiphytic conditions (log_2_FC: 2.0395 without temporal variation). Finally, genes encoding cellulose synthase (*bcsABC*) were uniquely associated with the epiphytic niche, with *bcsB* gene, a c-di-GMP-binding regulatory component of cellulose biosynthesis, significantly upregulated (log_2_FC: 2.159) relative to the apoplast. As cellulose is a major component of the extracellular matrix, its induction underscores the importance of biofilm-associated persistence during epiphytic colonization.

#### Type IV secretion system and DNA exchange

Genes encoding the type IV secretion system (T4SS) were strongly expressed during early epiphytic colonization. WGCNA placed *virB1, virB7–virB10*, *virC1,* and *virD4* in the epiphytic-specific green module, while *virB1, virB11,* and *virB4* clustered in the purple module associated with early infection in both niches. Plasmid-borne genes involved in conjugation, *trbD* and *trbB*, were significantly upregulated at 8h epiphytic growth (Table S1), and additional transfer genes (*trbCEFGIJKL*) were associated with epiphytic and early apoplastic networks (purple and midnightblue modules) but showed higher expression under epiphytic conditions. The transcriptional regulator *traJ* was also upregulated (log_2_FC: 1.4), suggesting that horizontal gene transfer is favored during early leaf-surface colonization. Genes involved in natural transformation including genes encoding competence proteins, ComEA, ComF, DprA (SMF, a DNA-processing protein), genes encoding SMG proteins involved in nonsense-mediated RNA decay (post-transcriptional regulation), were co-expressed. A gene encoding HU DNA-binding protein, that facilitates biofilm formation by anchoring extracellular DNA (eDNA) to bacterial cells, was co-expressed, indicating a potential role in stabilizing biofilm architecture and promoting genetic exchange on the leaf surface.

#### Cell-wall degrading enzymes

Genes encoding cell wall–degrading enzymes (CWDEs) detected primarily under epiphytic conditions included glycoside hydrolases (GH5, cellulase A; GH30; GH39/*xynB*), glycosyltransferases (GT2 and GT4 (E2P69_RS12875)), cellulose synthase (*bcsABC*), and rhamnogalacturonan lyase. Early epiphytic growth (8h) showed strong induction of a cellulase family protein (*E2P69_RS19010*), and two GH5 genes encoding endo-β-1,4-glucanases involved in cellulose breakdown (*E2P69_RS00220* and *E2P69_RS00235*), indicating active cellulose remodeling during surface colonization.

##### Stress Tolerance and Defense Against Environmental Challenges

Epiphytic cells expressed a broad stressome encompassing oxidative, osmotic, and envelope stress responses, along with multidrug efflux (*emrA*, *emrB*, up to 3.7-fold, at 8h) and an AlgE-family outer membrane protein (putative exopolysaccharide export).

#### Oxidative stress responses

Key regulators included *ohr* and *ohrR* (organic hydroperoxide resistance), and a MarR family transcriptional regulator, involved in detoxifying oxidative byproducts. Catalase *katB,* involved in protecting cells from hydrogen peroxide toxicity was induced (log_2_FC: 2.99). Unlike *Pseudomonas syringae* [22], which preferentially induces oxidative defenses in the apoplast, genes encoding oxidative stress enzymes (*ahpF, ahpC, ohr, katB,* and *sodA*) were more strongly induced on the leaf surface in *X. perforans.* This contrast highlights a fundamental difference in strategy: whereas *P. syringae* mounts a reactive oxidative stress response primarily after entering the apoplast, *X. perforans* activates this response early, suggesting either ROS pathway induction during initial colonization or anticipatory defenses on the leaf surface to withstand oxidative bursts before host entry. The OxyR regulon, comprising *oxyR*, *ahpCF*, *katA*, *katB*, and *trxB* [41], was co-expressed, with higher transcript levels in epiphytic vs apoplastic growth (log_2_FC ∼2-4), indicating activation of hydrogen peroxide– inducible defenses. OxyR additionally regulates peroxiredoxin and thioredoxin reductase and has been implicated in T6SS regulation (Antar 2022). The *cyoABCD* operon, encoding cytochrome-o-ubiquinol oxidase involved in redox balance and oxidative stress adaptation, was detected in epiphytic conditions. Deletion of these genes in *X. oryzae* increases sensitivity to hydrogen peroxide and reduces virulence. Additional redox genes included *yneN* (thioredoxin) and *rbn* (UPF0761 YihY family inner membrane protein), and *gshB* (glutathione synthase), supporting glutathione-based detoxification. Green module also included gene encoding rubredoxin, iron-sulfur protein involved in electron transfer, reduction of superoxide, and metabolism.

#### Osmotic stress response

Epiphytic cells upregulated *treA* (trehalase, enabling trehalose utilization under high osmolarity), and the high-affinity ATP-driven K^+^ uptake system (*kdpABCD*), forming an ATP-dependent K⁺ pump that maintains intracellular potassium levels and intracellular turgor pressure under osmotic stress. Interestingly, this system was highly upregulated under epiphytic conditions irrespective of timepoints showing transcripts levels of about 4. Regulators such as *ompR* (response regulator, activated upon sensing of changes in osmolarity by membrane bound sensor kinase, EnvZ) and *osmC* (osmotic and peroxide stress protection) were also detected, indicating porin remodeling and peroxide/osmotic protection.

#### Envelope stress response and intermicrobial competition

Copper homeostasis genes (*copA*, *copB*, and copper chaperone) and *baeS* (envelope stress sensor) were expressed, alongside mercury resistance. Interestingly, *baeS* is indicated to be activated in response to cell envelope damage such as an attack from competitor’s T6SS (Koler et al. 2016). T6SS components were distributed across modules (*tssM* in magenta; *hcp* in green, and *vgrG*/ *vipA* in the greenyellow module), with *tssM* significantly higher in epiphytic conditions (log_2_FC: 1.23), consistent with competitive interactions on the leaf surface. The sigma-E regulon (*rpoE*, *rseA*, *mucD*), mediating adaptation to membrane-perturbing conditions [42], was unique to epiphytic niche.

##### Protein folding/misfolding and DNA repair

Green module genes included genes involved multiple DNA repair systems *radA* and *msrA* (peptide-methionine S-oxide reductase, which repairs oxidatively inactivated proteins), DNA repair photolyase, MiaB family radical SAM proteins, *fpg* (a bifunctional DNA-formamidopyrimidine glycosylase/DNA-(apurinic or apyrimidinic site) lyase), and the *msrPQ* system for repairing oxidized periplasmic proteins. Additionally, chaperones (*dnaK*, *grpE*, *hslO*) and proteases (*lon*,*degQ*) were co-expressed, preventing protein misfolding. Clp, an indirect regulator of GroEL/GroES, was also detected, along with phosphatidate cytidylyltransferase, which modifies membrane fatty acid composition under stress. DnaK and GrpE proteins prevent aggregation of stress-denatured proteins. DegQ family serine endoprotease, *mucD*, degrades transiently denatured and unfolded proteins which accumulate in the periplasm following stress conditions. Genes involved in protein quality control and redox regulation, such as *hslO* and thiol-disulfide interchange proteins (*dsbA/L*), were expressed, ensuring proper folding of periplasmic and virulence proteins. The *msrPQ* system, which is significantly upregulated at 8h, repairs oxidized periplasmic proteins containing methionine sulfoxide residues, protecting against oxidative stress, while *lon* endopeptidase selectively degrades mutant, or abnormal proteins. Other co-expressed genes include *hslO* chaperone that is redox regulated and protects thermally unfolded and oxidatively damaged proteins. STRING network analysis linked *ahp* and *yneN* to these stress-response pathways.

Genes encoding DNA recombination protein RmuC (anti-phage/anti-MGE), recombination regulator RecX, recombinase RecA, integration host factor alpha IhfA, were co-expressed in the green epiphytic module (typical log_2_FC ∼1.2, *ihfA/recA* 0.8 at 8h), consistent with IHF-mediated nucleoid remodeling that directly regulates promoters of type I-VI secretion system genes, as demonstrated in *Dickeya* [43]. Co-expression of *dsbA/L* suggests DsbA/L-dependent disulfide bond formation supporting folding of periplasmic virulence proteins and type III effectors during epiphytic colonization [44]. NUDIX hydrolases, which contain an hrp box and are therefore predicted HrpX targets, were expressed on the leaf surface together with NAD salvage genes, encoding nicotinamide-nucleotide adenylyltransferase and nicotinate phosphoribosyltransferase (log_2_FCs: 4.0815 and 3.2490, respectively), despite the absence of *hrpX* expression under epiphytic conditions. Stressosome-associated components, including *rsbR* and a STAS-domain protein, were also expressed, indicating activation of global stress signaling pathways during epiphytic colonization.

#### Metabolism

##### Amino acid metabolism

Metabolic pathways enriched epiphytically included aspartate 1-decarboxylase supporting β-alanine production for pantothenate and CoA biosynthesis, phospho-2-dehydro-3-deoxyheptonate aldolase initiating the shikimate pathway for aromatic amino acid synthesis, and phenylalanine degradation. Induction of *phhA* (log_2_FC: 2.3) and *hppD* (log_2_FC: 1.04) suggests active utilization of phenylalanine, a precursor in plant phenylpropanoid defense, converting it to tyrosine during colonization, and hints at phenylalanine degradation by *Xp* early during epiphytic colonization and before infection of apoplast, similar to that observed in *P. syringae*.

##### Fatty acid metabolism

Genes *fadL* and *fadB*, involved in long-chain fatty acid uptake and b oxidation, were co-expressed under epiphytic conditions. Co-expression of the gene encoding the lipid-binding protein, DegV, further suggests coordinated regulation of fatty acid storage and catabolism, potentially restricting fatty acid export via membrane vesicles and linking lipid metabolism to vesicle regulation in the epiphytic niche [45].

#### Overcoming nutrient-limiting conditions

Genes encoding at least nine TonB-dependent receptors (TBDRs), three energy transducers, and *exbD4* transporter were co-expressed under epiphytic conditions, indicating active nutrient acquisition. Early epiphytic upregulation of the gene encoding ferric-rhodotorulic acid/ferric-coprogen receptor FhuE, and phosphate-selective porins (*oprO/oprP*) suggests iron and phosphate limitation on the leaf surface. Co-expression of iron homeostasis genes (those encoding iron-sulfur cluster assembly protein SufT, ferredoxins (FdxA), iron-sulfur proteins (IscA), and accessory proteins such as HesB (IscA family), and Grx4 family monothiol glutaredoxin (GrxD) involved in iron-sulfur cluster biogenesis and sensing iron starvation, further support persistent iron-depleted conditions throughout epiphytic colonization.

#### Adaptation to plant defense metabolites and stress responses

Plants deploy antimicrobial metabolites during pathogen attack, including volatile esters and aldehydes. During epiphytic colonization, *Xp* co-expressed the propionate catabolic regulator *prpR*, suggesting active metabolism of plant-derived propionate during epiphytic colonization, potentially counteracting stomatal closure and maintaining access to entry points [46]. In addition to esters, plants emit volatile organic compounds (VOCs) such as methanol, which can lead to the formation of formaldehyde—a toxic intermediate that must be detoxified. Concurrently, formaldehyde detoxification genes (*frmR*, *frmA*, *gfa*, and *fghA*), including *gfa*, encoding 5-hydroxymethyl glutathione synthase, and an aryl aldehyde reductase (E2P69_RS15385), were strongly induced during epiphytic conditions, consistent with active detoxification of aldehydes and a robust glutathione-dependent system for mitigating aldehyde and oxidative stress on the leaf surface [47,48].

Eight genes encoding type II toxin–antitoxin (TA) system components were significantly overexpressed during epiphytic (avg. log_2_FC: ∼1.5) growth relative to apoplastic conditions, with four induced specifically during early epiphytic colonization and remaining four throughout the epiphytic phase. TA systems are well established stress response modules that promote bacterial persistence (persister state) under nutrient limitation and oxidative stress by inducing reversible growth arrest or stabilizing essential pathways [49–51]. These findings highlight how *X. perforans* integrates metabolic flexibility, detoxification mechanisms, and stress-adaptive strategies to thrive in the chemically challenging epiphytic niche.

### Early host contact primes horizontal gene transfer and effector deployment

The purple module was strongly associated with early infection following initial host contact, independent of niche. Of its 136 genes, 44 were significantly upregulated during early epiphytic colonization (8h vs 24h), whereas 124 were induced during early apoplastic infection (24h vs 72h). This module included plasmid-borne genes involved in surface attachment, horizontal gene transfer, metabolism, and virulence, notably type IVa pilin genes (*pilO, pilP, pilQ*), T4SS components (*virB1, virB11*), and conjugation genes (*trbC, trbE, trbF, trbG, trbI, trbJ, trbL*, and recombinase), all highly expressed at early time points. Plasmid-borne effectors (TAL effector, *avrHah1* and *xopH*) were co-induced with chromosomal effectors (*xopQ* and *xopE*), athough core T3SS genes were absent from this module. While *avrHah1* expression showed significant induction during early epiphytic growth, other effector transcripts were predominantly upregulated during early apoplastic infection. Central to this early transcriptional program was the global post-transcriptional regulator, *csrA,* which controls many different physiological processes and virulence associated traits [52–54], and is present as both chromosomal and plasmid-borne copies in AL65. Both copies were induced during early epiphytic growth, whereas only the plasmid-encoded *csrA* was upregulated during early apoplastic colonization, underscoring its likely role in coordinating niche-specific regulation of virulence, metabolism and stress adaptation [27,55]. Additional co-expressed genes supported this early adaptive state in apoplast, including those involved in amino acid and nucleotide metabolism (*ggt, trpE, glyA, aroG*) and stress resistance (peroxiredoxin, alcohol dehydrogenase, and multidrug MFS transporters). This module also contained *acnB* (aconitase B), a [Fe–S] enzyme involved in both conversion of citrate to isocitrate in the TCA cycle flux and redox/iron sensing [56,57]. These observations are consistent with the view that aconitases integrate metabolic demands with redox and iron homeostasis [58,59]. Given the abundance of organic acids such as citrate in the tomato apoplast [60,61], induction of citrate transporter (*citM***) (**log_2_FC = 5.03) in the early apoplast is consistent with a metabolic program that leverages apoplastic citrate during early colonization as previously reported [62,63]. Thus, early induction of *acnB* likely integrates metabolic readiness with stress sensing during initial host engagement.

### *Xp* is equipped with unique adaptive strategies for colonizing and multiplying within the apoplastic environment that include chemotactic sensing, tolerance to nutrient limitation, active cell division and biogenesis, secretion of cell wall–degrading enzymes via the Type II secretion system, deployment of the Type III secretion system, and multiple stress-response pathways to counteract reactive oxygen species (ROS)

In the apoplastic niche, the yellow and greenyellow modules were niche-specific, together comprising 430 genes, of which 299 were significantly upregulated relative to epiphytic samples. In contrast to the temporal stability observed during epiphytic colonization, apoplastic infection was strongly structured over time. Early apoplastic infection was dominated by the blue and midnight blue modules, in which 575 of 629 genes were significantly upregulated at 24h compared with 72h. Conversely, the black module was specific to late apoplastic infection containing 311 genes, of which 308 were significantly upregulated at 72h compared to 24h. *Chemosensing, chemotaxis and quorum sensing*. Early apoplastic infection (blue and midnight blue modules) consists of genes involved in chemotaxis and signal transduction, including *raxH* and *cheW*. RaxH, a HAMP-domain histidine kinase, forms the RaxH/RaxR two-component system, which senses host-derived signals and regulates virulence-associated pathways. In *Xanthomonas oryzae*, this system controls *raxABC* Type I secretion system, critical for virulence [64]. Early apoplastic colonization was also marked by induction of *guaA*, involved in nucleotide biosynthesis, and two EAL-domain phosphodiesterases (E2P69_RS08940 and E2P69_RS13670), suggesting active modulation of c-di-GMP levels. Consistent with this, c-di-GMP-responsive regulator Clp was active during early apoplast infection but was significantly downregulated by 72h. Reduced c-di-GMP levels activates Clp, which in turn promotes expression of virulence genes [65]. Additional apoplast-specific sensory systems included PAS domain histidine kinases (E2P69_RS07440 and E2P69_RS07460) and HAMP-domain kinase *colS*, orthologous to the *Pseudomonas* ColS/ColR system involved in metal sensing, and low pH adaptation [66–68]. The *ravR* from RavS/RavR two-component system, identified in the apoplast-associated yellow module, is an EAL domain phosphodiesterase, modulates the swimming to virulence transition through c-di-GMP hydrolysis [69].

Late apoplastic infection featured expression of additional sensory kinases, including GAF-domain histidine kinase (E2P69_RS12270), two HAMP domain-containing histidine kinases (E2P69_RS01145 and E2P69_RS13405) and two PAS domain-containing histidine kinases (E2P69_RS04400 and E2P69_RS04625), suggesting continued environmental sensing as infection progresses. Overall, enrichment of EAL-domain enzymes in apoplast-specific modules indicates sustained c-di-GMP degradation during endophytic growth, whereas GGDEF-domain proteins associated with epiphytic modules suggest that c-di-GMP synthesis predominates during leaf-surface colonization.

#### Overcoming the nutrient limitation

Apoplastic niche-specific genes included those associated with nutrient limitation, suggesting that the pathogen experiences restricted nutrient availability, either due to host strategies aimed at nutrient sequestration or as part of the plant’s defense response.

##### Phosphate limitation

*Xp* senses phosphate limitation as early as 24h during apoplastic colonization and maintains phosphate acquisition and signaling throughout infection. Early apoplast samples showed induction of phosphate selective outer membrane porins, *oprO/oprP* (E2P69_RS01730, E2P69_RS15350), mediating uptake of orthophosphate and polyphosphate under starvation [70–72]. High-affinity phosphate import was further reflected by co-expression of the *pstSCAB* ABC transporters dedicated to high-affinity inorganic phosphate (P_i_) uptake [73,74] in both early and apoplast-specific modules, together with *pitA* (low-affinity P_i_ importer) and *ppx* (exopolyphosphatase that degrades polyphosphate to orthophosphate), indicating remodeling of P_i_ uptake and intracellular polyphosphate turnover in response to limitation [75]. By 72h, late apoplast samples showed co-expression of the phosphate-starvation regulators *phoR/phoB* and polyphosphate kinase (*ppk*), consistent with a shift from immediate uptake to regulatory and storage-based strategies characteristic of prolonged phosphate stress [76]. Recent work emphasizes how cytoplasmic phosphate levels and PstSCAB flux jointly tune Pho activation, preventing inappropriate responses while enabling adaptation to P_i_ deprivation [73,77]. Activation of the Pho regulon at this stage suggests that *Xp* manages its fitness costs when P_i_ is locally sufficient [74] and activates it later during disease progression while adapting to host mediated phosphate sequestration.

##### Sulfur metabolism

Pathogens rely on sulfur from their plant hosts and can manipulate plant sulfur transport systems to acquire sulfur during infection, while plants interpret sulfur depletion as a signal of pathogen invasion and intensify sulfur sequestration as a defense mechanism. This nutrient tug-of-war is critical because sulfur is essential for antioxidant defenses and redox homeostasis. While plants primarily assimilate inorganic sulfur as sulfate, bacteria can utilize both inorganic sulfate and organic sulfur such as sulfonates (secondary source under limiting conditions) and sulfate esters. In our dataset, sulfur metabolism genes are largely enriched in the apoplastic niche, with the exception of a single SulP family inorganic anion transporter for sulfate uptake expressed epiphytically, contrasting with the predominantly epiphytic sulfur-uptake profile reported for *P. syringae* [22]. Apoplast-specific modules revealed coordinated sulfur acquisition and assimilation programs. The yellow module contained *cysA* (sulfate/molybdate ABC transporter), while the early apoplast blue module included sulfate ABC transporter permeases (*cysW* and *cysT*), sulfite reductase (*cysJIH,* log_2_FC: ∼5), sulfonate ABC transporters (log_2_FC: ∼5), and taurine catabolism genes (*tauA* and *tauD;* log_2_FC: 5.17 and 3.75). The midnight blue module featured the sulfate-starvation regulator *cysB*, and sulfate adenylyltransferase (*cysC* and *cysD*), all showing strong induction at 24h relative to 72h. Because *cysB* activates sulfate starvation responses, including *tau* operon, this co-expression pattern suggests early sensing of sulfur limitation and a capacity to switch from sulfate to organic sulfur sources as stress intensifies. Similar induction of sulfonate and taurine utilization genes under PAMP triggered immunity has been reported previously, consistent with host-mediated sulfur sequestration or oxidative damage to Fe-S cofactors producing sulfur-starvation like responses [78].

##### Nitrogen metabolism

Genes associated with nitrogen metabolism were strongly induced during early apoplastic colonization, showing >2 log_2_FC higher expression relative to epiphytic and late apoplastic conditions. These included nitrate assimilation genes (*nasD, nasE*), nitrate reductase (E2P69_RS17555). Additional components, including an ammonium transporter (*amtB*), the P-II family nitrogen regulator (*glnB*), and glutamate synthase (*gltD*) were also upregulated at 24h post-inoculation. This coordinated induction indicates active nitrate assimilation upon entry into apoplast, likely reflecting nitrogen limitation imposed by host defenses. Similar niche-specific nitrate utilization has been reported in *P. syringae,* where nitrate reductase expression was detected exclusively in cells grown in apoplastic wash fluid, but not in macerated tissue, indicating nitrate utilization is a niche-specific adaptation [79]. Presence of localized nitrate pockets on leaf surfaces were demonstrated using bioreporter experiments, although carbon remains the primary limiting factor, followed by nitrogen [80]. These observations suggest that coordinated strategy for nitrogen assimilation and incorporation into amino acid biosynthesis during early infection, as host defenses often restrict nitrogen availability as part of immune responses.

##### Amino acid metabolism

Amino acid metabolism and translational capacity were actively remodeled during early apoplastic colonization. The blue module showed induction of phenylalanine-tRNA ligase subunits (*pheS/pheT*), consistent with a surge in translational demand to support rapid growth and the synthesis of virulence and stress-response proteins. Despite reported glutamate abundance in the tomato apoplast [81], *gltB* (glutamate synthase) was induced in the midnight-blue module, suggesting roles in ammonia assimilation, maintenance of central carbon-nitrogen balance, and redox control. Induction of *gltP* (glutamate/asparate symporter) and glutamate-5-kinase further links glutamate metabolism to chemotaxis and osmoprotection via proline biosynthesis [22]. Expression of *thrE* (threonine/serine exporter) in blue module further suggests that *Xp* actively prevents toxic intracellular accumulation of amino acids while fine-tuning osmotic balance. Notably, unlike *Pseudomonas syringae*, which primarily upregulates amino acid transporters *in planta* [22], *Xp* engages both uptake and biosynthetic routes early in apoplastic colonization. This may suggest *Xp*’s strategy to prioritize metabolic autonomy, securing key amino acid precursors, maintaining redox-osmotic homeostasis, while also equipping itself with the translational capacity to fulfill the energy demands of virulence.

Genes involved in broader amino acid biosynthesis were distributed across apoplast-associated modules. The yellow module showed strong expression of the *metLB* operon and *metXS*, indicating activation of both canonical and alternative methionine biosynthesis pathways, similar to *Pseudomonas*. Branched-chain amino acid biosynthesis genes (*ilvM, ilvG, ilvC)* and shikimate pathway enzymes (chorismate mutase) were induced across apoplastic infection, while *trpC/trpD* for tryptophan biosynthesis, threonine (*thrABC*) along with *thrE* (a threonine/serine exporter for excess amino acids removal) and arginine (*argHCABEGF*) biosynthesis genes were primarily induced during early apoplast colonization. Early apoplastic colonization also supported proline catabolism, as indicated by expression of *putA*, encoding proline dehydrogenase, where it oxidizes proline to glutamate, providing carbon and nitrogen and linking nitrogen metabolism, osmoprotection, and T3SS induction [82]. Histidine biosynthetic genes (*hisA, hisB, hisC, hisD,* and *hisH*) were also induced, consistent with histidine limitation in the apoplast, and the importance of histidine homeostasis for virulence, redox balance, and metal stress tolerance [83–85].

##### Sugar metabolism

Early apoplastic colonization by *Xp* is characterized by strong induction of carbohydrate acquisition and exopolysaccharide (EPS) biosynthesis. The midnight blue module included *oprB* (formerly *rpfN*), a carbohydrate-selective porin essential for growth and virulence (log_2_FC: 1.50). Loss of this porin reduces sugar uptake and induces compensatory cell wall–degrading enzymes (CWDEs) to liberate plant-derived sugars, underscoring its role in nutrient acquisition [86]. Carbohydrates imported via *oprB* likely fuel EPS production, supported by co-expression of *gumEFGHIJ* in the early apoplast (log_2_FC: 1.20-2.47) and *gumKLM* in the yellow module (log_2_FC: 2), suggesting that EPS production begins early and persists throughout colonization, contributing to biofilm formation and protection against host defenses. Additional sugar metabolism genes induced early included *scrK* (fructokinase), *fucP* (fucose permease), and sucrose utilization genes within the *sux* operon (*suxA*, *suxC*, and regulator *suxR*), consistent with the importance of sucrose uptake for virulence. Induction of succinyl-CoA synthetase (*sucC* and *sucD*), further links carbohydrate assimilation to TCA cycle activity and energy generation during apoplastic infection.

#### Other metabolism-related genes

Early apoplastic colonization was marked by induction of cobalamin (vitamin B_12_) biosynthesis genes (*cobU, cobQ*; log_2_FC: 2.57), supporting cofactor-dependent central metabolism and redox balance under nutrient limitation. Concurrent upregulation of the F_0_F_1_ ATP synthase (*atpAGDC*), and fatty acid biosynthesis genes (*fabFGDH*), indicates elevated energy investment and membrane remodeling including xanthomonadin pigment production, to maintain envelope integrity under host-induced oxidative stress during early apoplastic colonization. Early infection was further characterized by induction of ribosomal proteins, ribosome maturation factor (*rimP*), RNA polymerase subunits (*rpoBC*), and translation-associated factors, consistent with increased biosynthetic and virulence demands. In contrast, late apoplastic infection showed induction of the *moaEDCAB* operon, involved in molybdenum cofactor (Moco) biosynthesis. Enhanced Moco production likely activates molybdoenzymes required for redox regulation, anaerobic respiration, and nitrogen metabolism, facilitating metabolic flexibility and survival as the apoplast becomes increasingly oxygen and nutrient-limited [87].

#### Cell envelope biogenesis

Genes involved in peptidoglycan biosynthesis were strongly represented in the early apoplast-specific blue module, indicating large investment in robust cell envelope biogenesis. These include the *mur* gene cluster (*murB* through *murF*), which encodes enzymes responsible for sequential assembly of soluble peptidoglycan precursors, culminating in UDP-N-acetylmuramyl-pentapeptide synthesis [88]. Co-expression of multiple *fts* family cell division genes further indicates active cell wall remodeling and septation during early apoplastic infection.

#### Type II secretion system and cell wall degrading enzymes (CWDE)

Early apoplastic colonization was characterized by strong induction of type II secretion system (T2SS) components and associated CWDEs. These included glycoside hydrolases (GH98, GH2, GH3, GH92, GH95, GH97, GH99), cellulases (*celD*), beta-galactosidases (*galA*, *bga2*), and glycosyltransferases (GT41, GT2), together with core T2SS machinery genes (*xpsFMND*), highlighting active export of plant cell wall degrading enzymes facilitating nutrient acquisition [27,89]. Additional CWDEs uniquely expressed early included GH5, GH28, GH31, GH35 (*galD*), GH43, GH95, GH125, and GH127 (log_2_FC: ∼1 to 5 compared to late apoplast). These CWDEs target diverse substrates, such as cellulose, hemicellulose, and pectin, enabling pathogen to exploit plant derived carbohydrates during infection. Other CWDEs expressed uniquely in apoplast include GH5 cellulase (*egl*), cellulase (*engXCA*), GH3 (*bglX*), and chitinase GH19. Core components of a second T2SS (*xcsH* and *xcsC,* and *xcsF*) were detected throughout the apoplastic infection. Interestingly, some T2SS-related genes and glycosyltransferases (e.g. GT21, GH15, GH30) were expressed during later stages of disease, suggesting a sustained role of T2SS in virulence and tissue maceration as infection progresses.

#### T3SS system genes and effectors

During early apoplastic colonization, genes associated with cell division were co-expressed with T3SS components, indicating coordinated investment in growth and virulence. T3SS genes showed strong induction in early apoplast (avg. log_2_FC ∼4 relative to late apoplast and ∼5 relative to epiphytic growth). Most core *hrp* cluster genes, including chaperone *hpaB*, together with effectors (*xopV, xopN, xopZ, xopX, xopF1, xopK,* and *avrXv4*), clustered in the blue module, whereas additional effectors (*avrBs2, xopL, xopAD, xopI, xopAE, xopF2*, and *xopAV*) were enriched in the early apoplast-specific midnight blue module. Notably, several effectors were associated with the purple module expressed in at 24h while others (*xopAV, avrXv3)* were associated with the apoplast-specific greenyellow module, indicating spatiotemporal regulation of effector deployment across infection stages [27,90]. Early effector transcription in the absence of *hrp* gene expression suggests that some effectors, particularly plasmid-borne ones, may be pre-synthesized and poised for rapid delivery once a functional T3SS is assembled, or may utilize alternative delivery routes, as reported for AvrBs3 homologs [91].

#### Virulence regulon

The HrpG* core regulon encompassing T3SS structural genes, effectors, and metabolic regulators (*pcaQ, virK,* and *vanK*) was expressed primarily in the early apoplast-associated blue module. Plant cell-wall degradation releases phenolic compounds, including vanillic acid, 4-hydroxybenzoic acid, hydroxycinnamic acid. VanK mediates their uptake, while PcaQ regulates their catabolism, enabling detoxification and use of aromatic intermediates as carbon sources [92]. Additional HrpG* core regulon members including *phoC* (acid phosphatase), and PAP2 superfamily, genes were enriched in apoplast-associated greenyellow module, whereas chorismate mutase was detected in the yellow module, highlighting layered niche-specific deployment of HrpG*-regulated functions during apoplastic colonization.

#### LPS biosynthesis

A strong induction of LPS biosynthesis genes was detected during early apoplastic colonization. These genes are located in a conserved cluster between *metB* and *etfA* (E2P69_RS05020-RS05085), encoding enzymes required for assembly of the LPS core and O-antigen [93]. In addition, the *rmlABCD* operon, responsible for synthesis of dTDP-L-rhamnose, the direct precursor for rhamnose moieties incorporated into the LPS core and O-antigen, was highly expressed. During late apoplastic infection, induction of LPS export genes (*lptG*, encoding ABC transporter permease) was observed. Together, LPS biosynthesis and export likely support outer-membrane integrity and immune evasion throughout apoplastic colonization, particularly during prolonged colonization under stress conditions [94].

#### Camouflaging and stress management strategies of a pathogen

##### Early apoplast infection

Early apoplastic colonization by *Xp* is marked by activation of detoxification, redox balancing, and stress adaptation pathways that enable survival in the chemically hostile apoplast. The midnight blue module included *ligA* and *ligB*, encoding dual role extradiol catechol dioxygenases that catalyze oxidative cleavage of substituted catechols [47], detoxifying antimicrobial phytoalexins, while generating aromatic intermediates for metabolism [95]. Early apoplast samples also showed strong induction of cytochrome c biogenesis genes (*ccmE, ccmG,* and *ccmF*), supporting their critical role in electron transport and oxidative stress tolerance. In parallel, genes from the *dsbE* family, responsible for introducing disulfide bonds into periplasmic proteins, were also co-expressed, ensuring proper folding of virulence factors and stress-response proteins under oxidative conditions [92]. Both ccm and dsb pathways have been implicated in tolerance to reactive oxygen species (ROS) and phenazines [96].

The stringent response was also activated early, as indicated by induction of *spoT*, encoding bifunctional (p)ppGpp synthetase/guanosine-3’,5’-bis(diphosphate) 3’-pyrophosphohydrolase. SpoT controls cellular levels of alarmone nucleotides (p)ppGpp and maintains the balance between synthesis and degradation of (p)ppGpp that is important in adaptation to the changing environment. This global regulatory pathway reprograms transcription under carbon and nitrogen limitation, reducing ribosomal RNA synthesis and prioritizing stress survival and virulence [97,98]. Defense against plant-secreted antimicrobials was evident through strong expression of NADH-quinone oxidoreductase genes (*nuoBCDEFGHIJLMN*) in the blue module, and *nuoA* in midnightblue module, which mitigates toxicity from plant-derived quinones, while sustaining respiratory metabolism [99,100]. Additional stress-related genes were detected, encoding BatD oxygen tolerance proteins, SmeC multidrug efflux protein, and YicC/YloC family endoribonuclease. Presence of glyoxalase/bleomycin resistance protein/dioxygenase superfamily genes expressed during early apoplast colonization suggests mechanisms for detoxifying redox-cycling compounds are in place to maintain redox balance [101].

##### Strategies specific to the apoplastic niche that are important throughout apoplastic colonization

Pyrroloquinoline quinone (PQQ) biosynthesis genes, (*pqqE* and its chaperone *pqqD*), were detected in the apoplast-specific modules. The induction of these genes suggests that *Xp* leverages PQQ-mediated electron transfer to stabilize redox homeostasis under stress conditions [102]. The glycine betaine/L-proline transporter *proP* was expressed in the yellow module, specific to the apoplastic niche, and showed significant induction at 24h compared to 8h during epiphytic colonization (log_2_FC: 2.72), indicating sustained responses to water limitation. The Suf pathway (*sufD*, *sufC*, and early induction of *sufE*) was activated to maintain iron homeostasis and repair oxidatively damaged Fe-S clusters.

##### Late apoplast infection

During late apoplastic infection, additional systems were engaged to sustain viability under low-oxygen and nutrient-limited conditions. These included cytochrome bd complex *(cioA,* and *cioB*, log_2_FCs: 3.10 and 3.14 respectively) that enhances resistance to toxic compounds such as metals, nitric oxide, sulfide, cyanide, hydrogen peroxide, and peroxynitrite, supporting survival under oxidative stress and low-oxygen conditions typical of the apoplast. The gene *tolB*, detected during late infection, plays a role in TonB-independent uptake and cell envelope stability. Genes encoding the ClpXP protease system and stringent starvation protein (*sspB*) were expressed during late infection, suggesting involvement in protein quality control and stress adaptation. Additionally, *glpR*, a glycerol metabolic repressor that responds to glycerol 3-phosphate (G3P), a crucial module signal of systemic acquired resistance, was expressed during late infection. G3P mediated relieving of the GlpR-dependent repression of the *glp* gene cluster and imposing of bistable growth pattern [103] suggests that *Xp* may sense and respond to plant-derived G3P, adjusting its metabolic state to optimize survival under stress. A CHASE-domain histidine kinase (*PcrK*) was detected, which senses plant cytokinins and initiates signaling cascades that influence bacterial gene expression and help *Xanthomonas* adapt to the plant environment by altering virulence and improving resistance to oxidative stress [104]. Genes encoding polyhydroxyalkanoate (PHA) granule-associated proteins were induced, suggesting that when carbon sources are abundant, but nitrogen is limited, the pathogen stores excess carbon as PHAs. These granules may stabilize membranes under hypertonic conditions, aiding survival during plasmolysis. Additionally, osmoprotectant systems were activated, including, *yehX/yehZ*, glycine betaine transporters for osmotic stress tolerance, *egtB* involved in ergothioneine biosynthesis, and alpha-trehalose phosphate synthase, supporting trehalose mediated osmoprotection. Genes encoding cupin-domain protein, phosphomannose isomerase, was expressed alongside sterol-binding proteins and regulators of ubiquinone biosynthesis (protein kinase regulator of UbiI) that contribute to LPS assembly and membrane integrity, essential for virulence and phage resistance [105,106]. The apoplast undergoes an oxidative burst during infection, driven by plant defenses. The oxidation of apoplastic ascorbic acid by ascorbate oxidase alerts redox signaling, and its conversion to oxalic acid may further restrict pathogen growth. Genes encoding *sodC1* (superoxide dismutase) and catalases *(katE*, Mn-catalase), are expressed to neutralize ROS, and withstand oxidative stress. Interestingly, these oxidative stress response genes differ from those expressed in the epiphytic niche, indicating niche-specific compartmentalization of stress adaptation strategies in *Xp*.

### Shared motility and stress adaptation gene expression between late apoplastic and epiphytic phases indicates a transitional stage for pathogen dissemination and surface recolonization

The magenta module was associated with late apoplast infection and both epiphytic time points. Of the 341 genes, 328 were significantly upregulated during late apoplast infection relative to early infection, whereas only 69 genes showed significant differential expression between epiphytic time points, indicating largely similar expression profiles across epiphytic conditions. This module contained a large repertoire of motility and chemotaxis genes, including a complete flagellar gene cluster (E2P69_RS17025 through RS17080; RS17170 to RS17260) and multiple chemosensory components (*cheY, cheR2, cheD, cheB, cheY, cheZ, cheA, cheW, cheA2*), suggesting a coordinated re-activation of swimming motility and chemotactic navigation during late apoplastic and early epiphytic infection. This pattern mirrors the hierarchical architecture of flagellar biogenesis described in *Xanthomonas campestris*, where RpoN1 (σ⁵⁴) and FleQ initiate expression of *FT3SS*, rod, and hook genes, followed by FliA (σ²⁸) for filament assembly, with FlhA/FlhB/FlgM gating export and sigmafactor activity. Perturbation of FlhA/FlhB reduces infection, underscoring the virulence link [107]. Co-expression of *cheY* supports re-establishment of directional switching and chemotactic bias during late apoplast navigation [108]. The magenta module also encoded multiple c-di-GMP signaling components, including four EAL-domain proteins (E2P69_RS09020, RS17280, RS17845, E2P69_RS20925), five GGDEF-domain diguanylate cyclases (RS10540, RS10550, RS18400, (RS22020 & RS22030)), and a single *PilZ*-domain effector (RS17160) located near flagellar genes, indicating fine-tuning of motility and biofilm dispersal in response to intracellular c-di-GMMP levels. Additional sensory systems include three PAS-domain proteins (RS02915, RS11650, RS12280), two HAMP-domain proteins (RS17605, RS19020), and two STAS-domain proteins (RS20270, RS22520), which integrate environmental cues into motility regulation. Another oxygen and stress sensing histidine kinase induced in this phase was a PAS-domain hybrid histidine kinase homologous to StoS (XOO_0635 in *X. oryzae*), an O₂ sensor, whose deletion reduces tolerance to high osmolarity, sodium, and H₂O₂ and diminishes *inplanta* fitness [109]. Other stress adaptation genes expressed in this module include glutathione-s-transferase, glutaredoxin, and a glutathione-dependent formaldehyde-activating enzyme, which collectively support redox homeostasis. Antioxidant defenses are further reinforced by expression of *srpA* (catalase family peroxidase) and genes encoding carboxymuconolactone decarboxylase family protein, which is an antioxidant protein with alkyl hydroperoxidase activity. Components required for detoxifying reactive oxygen species include AhpC-associated systems, which depend on thiol reduction for regeneration, SodM (superoxide dismutase), and thioredoxin (TrxA), ensuring protection against oxidative damage and maintenance of protein integrity.

## Conclusion

This study revealed remarkably distinct patterns of RNA transcript expression both spatially and temporally for *Xanthomonas perforans* infection on tomato. This not only provides clues about the conditions that the pathogen encounters both epiphytically and apoplastically but also demonstrates the importance of these dual lifestyles and the ability to rapidly transition between lifestyles to successfully colonize the leaf, cause disease, and be able to disseminate. Here we present a coherent model that summarizes the niche-specific and infection-stage-specific gene expression patterns **(Fig 4)**. The early epiphytic lifestyle is characterized by chemosensing, motility, including both flagellar and twitching motility, uptake of scarce nutrients such as iron and phosphate, production of DSF and cellulose, conjugation, natural competence, and DNA repair. The overall epiphytic lifestyle also includes upregulation of genes to counter oxidative stress and repair or degrade damaged proteins, reduce osmotic stress through potassium pumps, synthesize cellulose, type II toxin-antitoxin systems, copper resistance proteins, the Type VI Secretion System, and genes suggesting a greater focus on cyclic-di-GMP synthesis compared to degradation. Alternatively, in the early apoplast the pathogen upregulates genes for reproduction or cell turn-over, including nitrogen uptake and assimilation, amino acid synthesis, RNA polymerase, ribosomes, carbohydrate uptake, and proteins for cell division organization and peptidoglycan synthesis while favoring genes for degradation of cyclic-di-GMP compared to synthesis. Additionally, the early apoplastic lifestyle is characterized by virulence traits, T2SS, CWDE, and T3SS and effectors while also showing signs of defense from the plant’s immune response including increased phenol uptake and degradation, quinone detoxification, the synthesis of fatty acids, xanthan, and LPS, in addition to the upregulated uptake of phosphate and sulfur to counter the plant’s sequestration of nutrients in response to infection. The late apoplast is defined by upregulated genes for molybdenum cofactor synthesis, for ROS detoxification such as catalase, and increased transmembrane transport of LPS. Further, WGCNA analysis revealed unique patterns of gene expression with modules specific to early colonization regardless of location, revealing co-expression of genes for type 4 pili, conjugation, and plasmid-borne effectors. Another module showed co-expression of flagella and chemotaxis genes for epiphytic and late apoplastic potentially showing these functions are of particular importance for the transition phase from the apoplast to the epiphytic surface for dissemination. Overall, the majority of epiphytic and apoplastic traits in *Xp* important for tomato colonization closely mirrored those observed in *P. syringae* on bean leaves. Notable differences, however, were evident in niche-specific metabolic and stress response strategies. In *Xp*, sulfur metabolism-related genes were co-expressed upon entry into the apoplastic niche, in contrast to *P. syringae*, where they were predominantly upregulated in the epiphytic niche. Additionally, oxidative stress-related genes in *P. syringae* were induced mainly after transition into the apoplast, whereas *Xp* exhibited distinct and non-overlapping sets of oxidative stress-related genes between the epiphytic and apoplastic niches. While this study reveals information into the different traits expressed during the lifecycle of this devasting bacterial spot pathogen, there is still more knowledge yet to be gained about the triggers and pathways for the initialization of the transition between these life stages to elucidate the full picture of this pathogen’s ability to infect its host.

**Fig 4.**
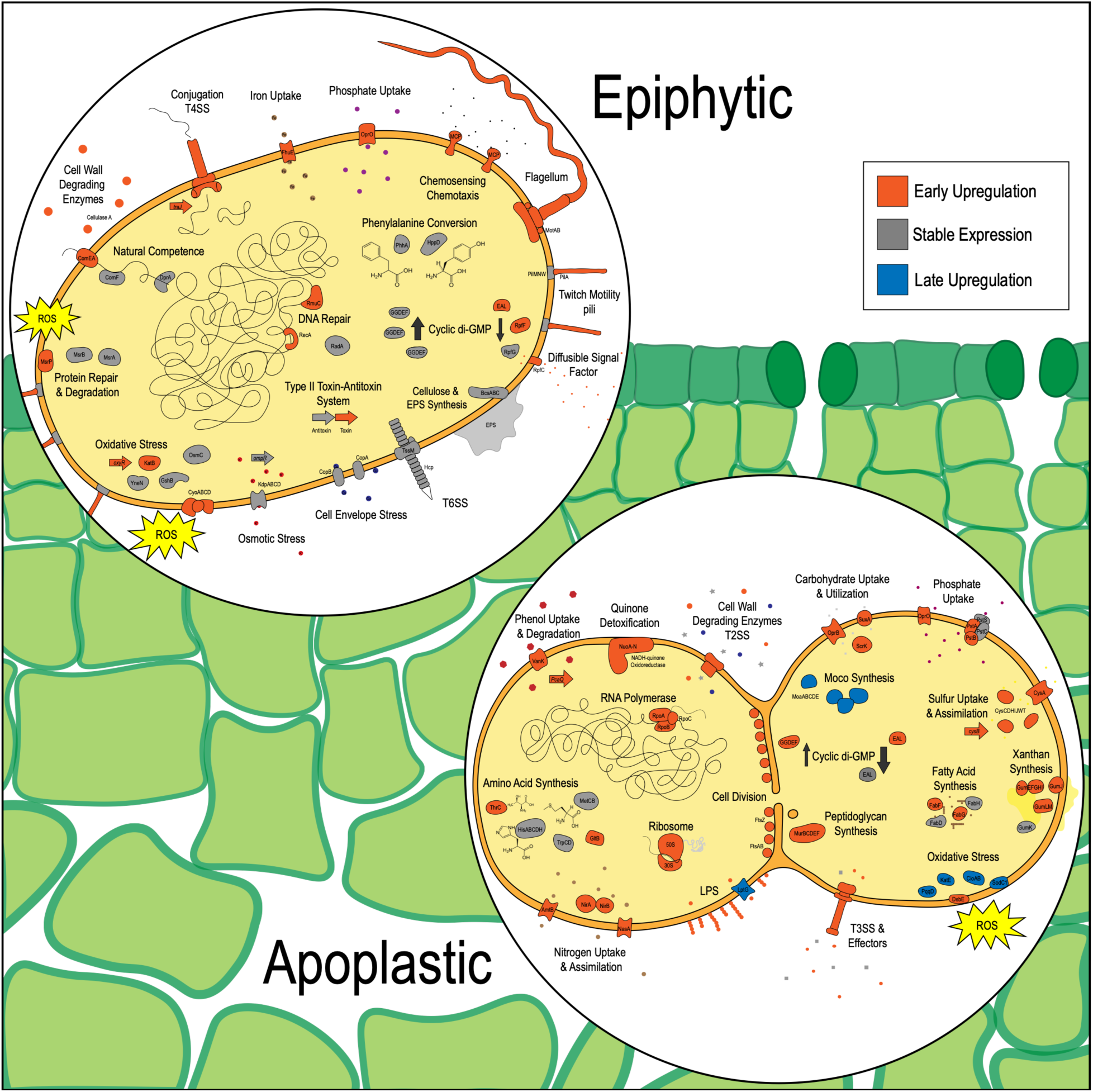
A summary of cellular systems upregulated only in epiphytic or apoplastic conditions. Note that not all upregulated systems and genes are shown. Components experiencing significant upregulation according to DESeq in the early timepoint (8 hours for epiphytic and 24 hours for apoplastic) are colored in orange, significant upregulation in late timepoint (24 hours for epiphytic and 72 hours for apoplastic) are colored in blue, and components that were not significantly different between timepoints are shown in grey.

## Data Availability

All samples sequence read archive are available on NCBI under the BioProject accession number PRJNA1405621. All codes and output files from different analyses conducted in this manuscript are uploaded on GitHub (https://github.com/Potnislab/Pathogen_inplanta-transcriptome.git).

## Supporting Information

Supplementary Tables

**S1 Table.** Host-specific depletion probes for enriching bacterial mRNA

**S2 Table.** Differentially expression genes categorized using KEGG pathway enrichment analysis

**S3 Table.** List of all the genes significantly associated within each module

**S4 Table.** Genes list grouped according to different systems within each condition (as referred in the text)

## Acknowledgements

We thank the Alabama Supercomputing Authority for granting access to their high-performance computing platform. We acknowledge Jessica Armstrong, Plant Science Research Center staff, and Alabama Ag Experiment Station for their support in conducting greenhouse experiments.

## Competing interests

The authors have declared that no competing interests exist.

## Notes

https://github.com/Potnislab/Pathogen_inplanta-transcriptome

### Competing Interest Statement

The authors have declared no competing interest.

### Funding

- U.S. National Science Foundation: 1942956

